## Supplementary Materials for "Landscape of allele-specific transcription factor binding in the human genome"

### Contents

Supplementary Figures S1-S12

Supplementary Tables S1-S5

Supplementary Methods

### Supplementary Figures

**Supplementary Fig. S1. The number of processed data sets (A, B) and the total number of filtered SNP calls (C, D) for different transcription factors and cell types (X-axis).**

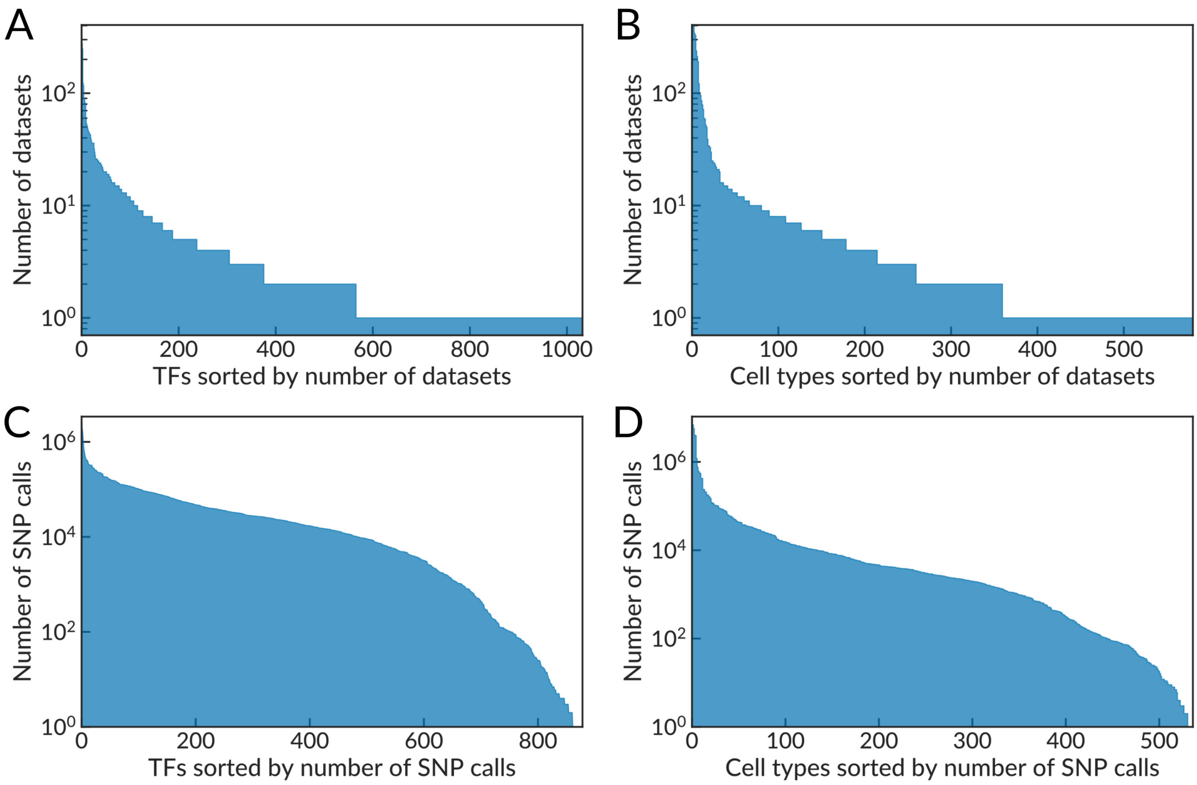

#### Supplementary Fig. S2. Results of the genome-wide BAD calling from ENCODE K562 data.

BAD calling applied to variant calls detected in K562 ENCODE data (ENCBS725WFV), each plot shows a single chromosome (note only a few false positive SNV calls on chrX and chrY). X-axis: chromosome position, bp. Y-axis: the allelic disbalance of individual SNVs. Horizontal green lines (ground-level of the plots) indicate results of the initial stage of the algorithm: the detection of SNV-free regions including deletions, telomeric, and centromeric regions. Horizontal light-blue lines: predicted BAD. Orange dashes: 'ground truth' BAD according to the COSMIC data (when available).

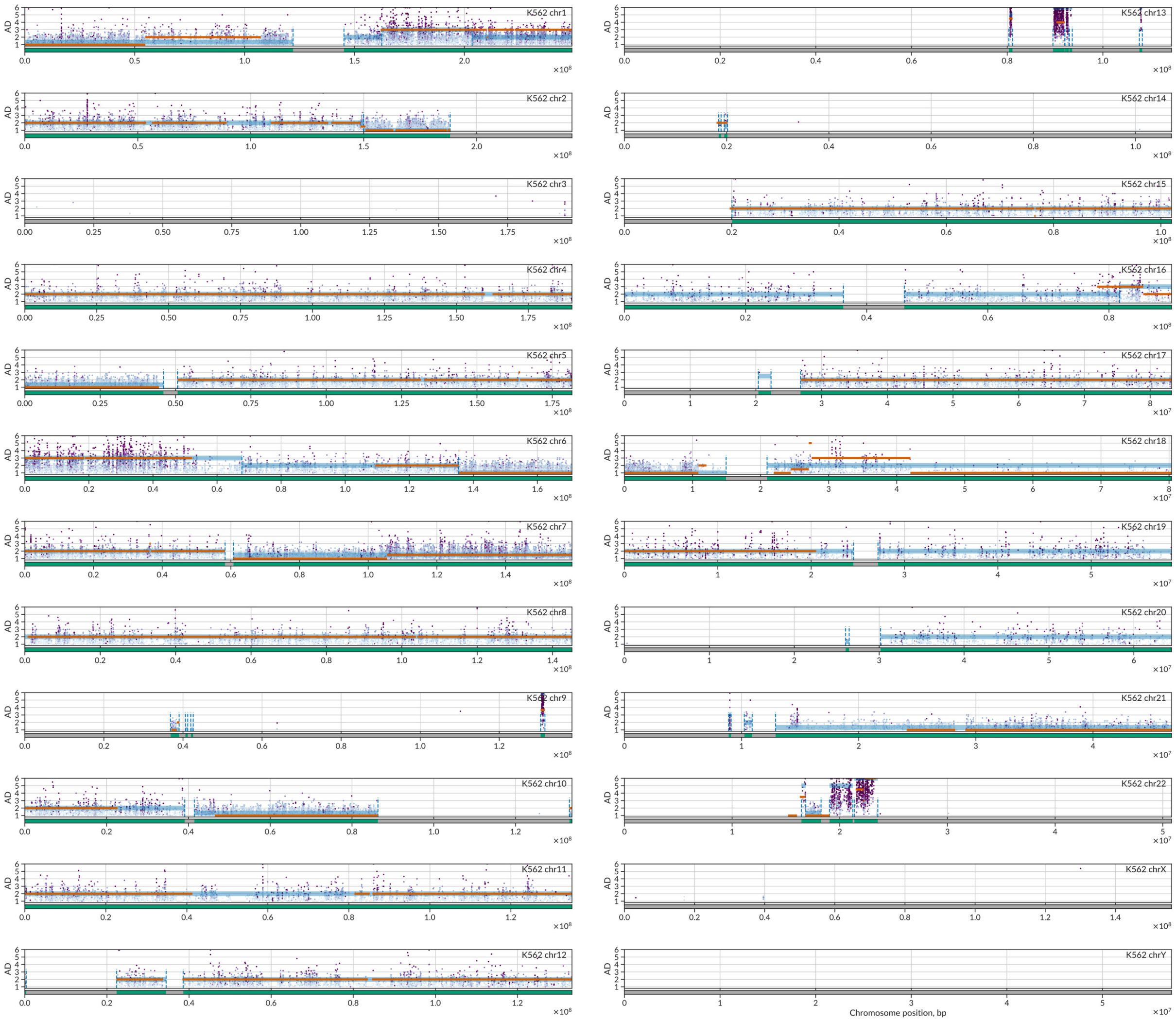

#### Supplementary Fig. S3. BAD prediction is successful for the most widespread BADs with the reliability comparable to that of the independent microarray data.

**(A)** X-axis: SNV-level Kendall $\tau_b$ rank correlation between predicted BAD and 'ground truth' BAD based on COSMIC data. Y-axis: SNV-level Kendall $\tau_b$ between total copy numbers estimated from the aCGH (Microarray-based Comparative Genomic Hybridization) and COSMIC data. For each SNV, the total copy number was estimated from the closest microarray probe.
**(B)** Bubble-plot illustrates the number of SNVs (bubble size) with particular predicted BAD (Y-axis) against the COSMIC data as the ground truth (X-axis). The most common BAD=2 is predicted correctly for the majority of SNPs.

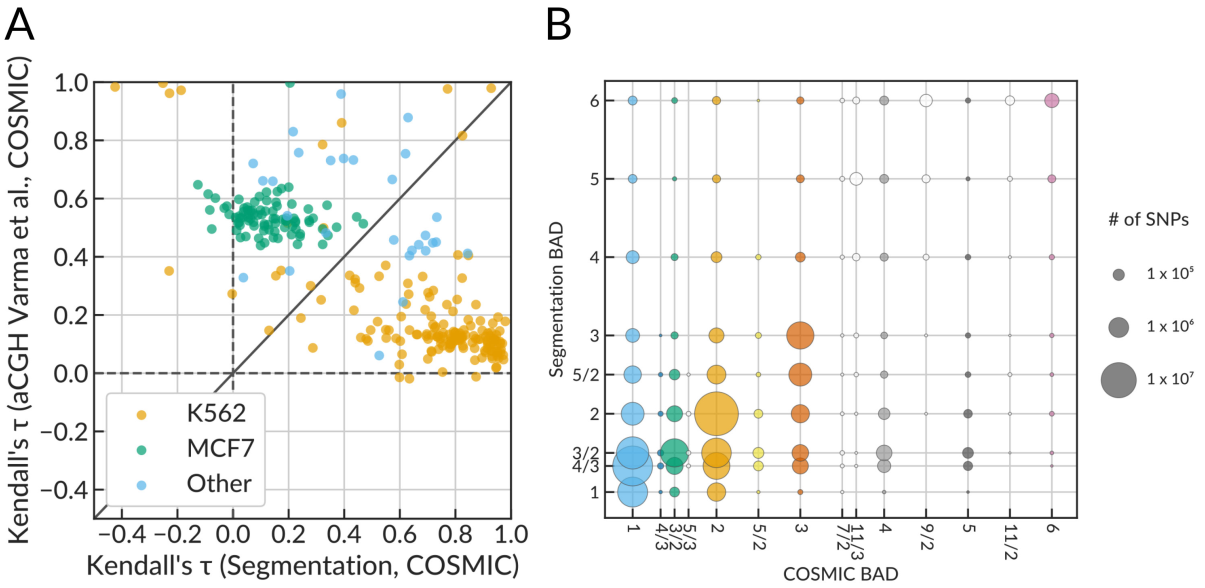


#### Supplementary Fig. S4. Distribution of distances between neighboring ASBs reveals ASB clustering at the scale of regulatory regions.

X-axis: the distance between neighboring SNVs, base pairs, $\log_{10}$. Y-axis: distribution density. SNPs with significant ASB events (FDR-corrected $P < 0.05$) obtained by **(A)** TF-aggregation and **(B)** Cell type-aggregation are shown in orange. SNPs with non-significant ASBs (non-ASB) are shown in gray.

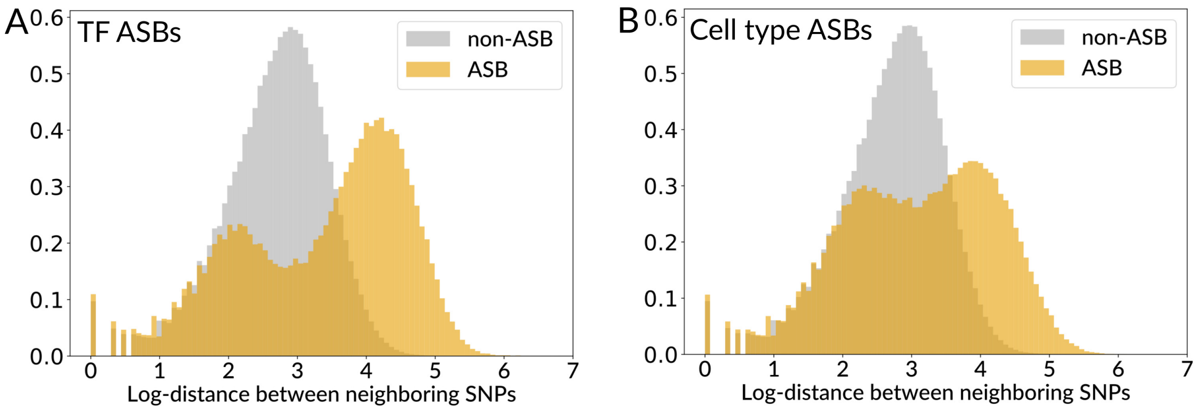

**Supplementary Fig. S5. ADASTRA ASB-SNPs overlap those found in other ASB collections, allele-specific DNase-accessibility sites, and raQTLs.**

**(A)** Venn diagrams illustrating the overlap between existing ASB collections and ADASTRA data.
**(B)** The number (top subpanel) and the fraction (bottom subpanel) of SNPs (dbSNP IDs) with candidate ASBs overlapping allele-specific DNase accessibility sites (imbalanced DNase sites) and not imbalanced DNase sites. X-axis: ASB significance threshold for TF-ASBs (-$\log_{10}$FDR-corrected P-value). The default threshold of 0.05 is shown as a dashed line.
**(C-D)** The number (top subpanels) and the fraction (bottom subpanels) of SNPs with candidate ASBs overlapping raQTLs in K562 **(C)** and HepG2 **(D)**, ASBs of the respective cell types were used in the comparison. X-axis as in **(B)**.

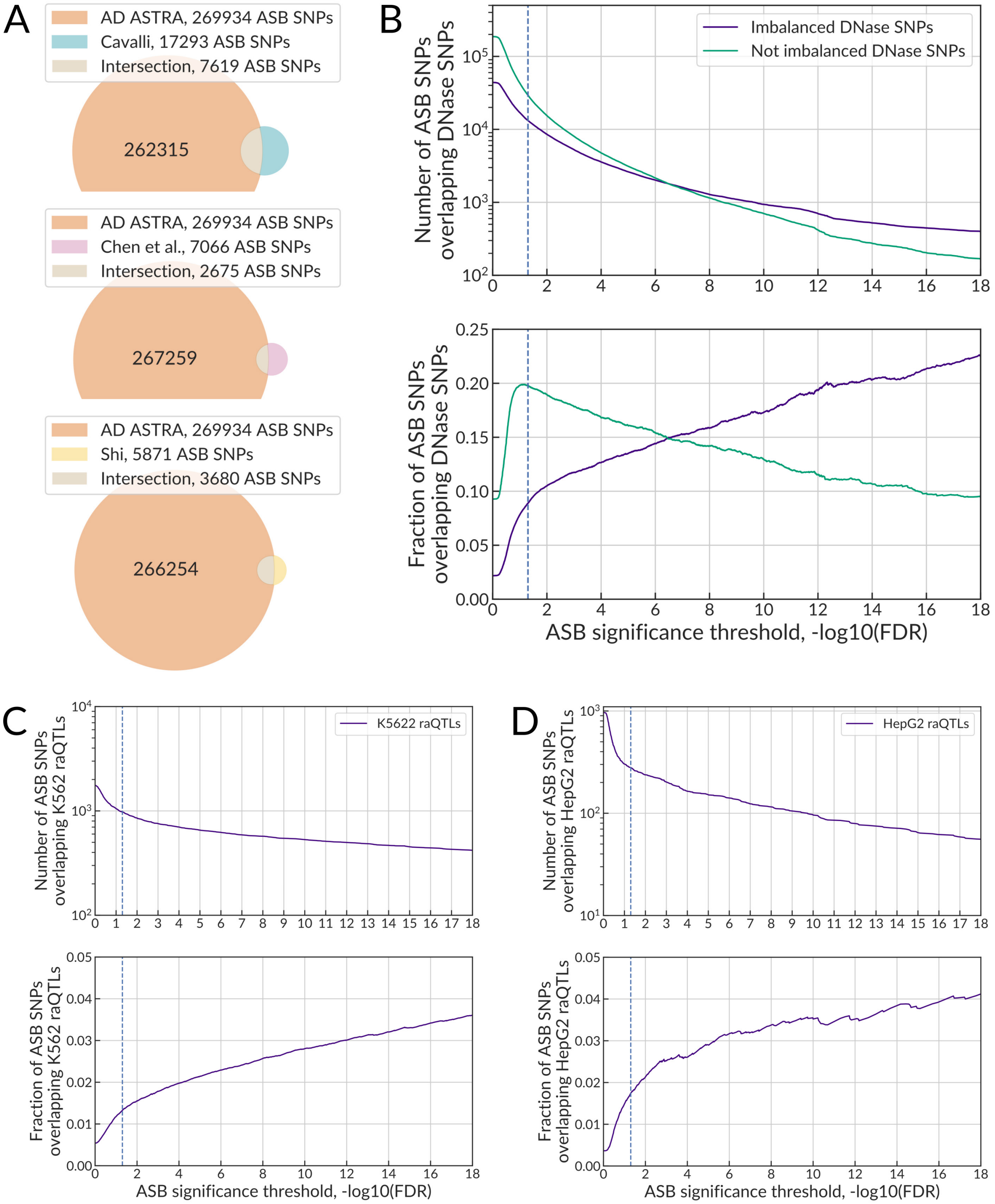

#### Supplementary Fig. S6. Particular TF pairs exhibit a preference for co-ASBs.

Cluster map of co-ASBs for TF-TF pairs with >100 co-ASBs. Each cell represents a TF-TF pair with SNVs exhibiting ASB events for both TFs. The coloring scale shows significance (-log$_{10}$P-value, FDR-corrected for multiple tested TF-TF pairs), the enrichment was estimated with Fisher's exact test. All significant ASBs were used as the background set. Non-significant enrichment is shown in white. Orange frames denote known TF-TF interactions with the combined score > 0.7 according to the STRING-db. The plot was produced with the *clustermap* of the seaborn Python package with the *cosine* similarity and *average* clustering.

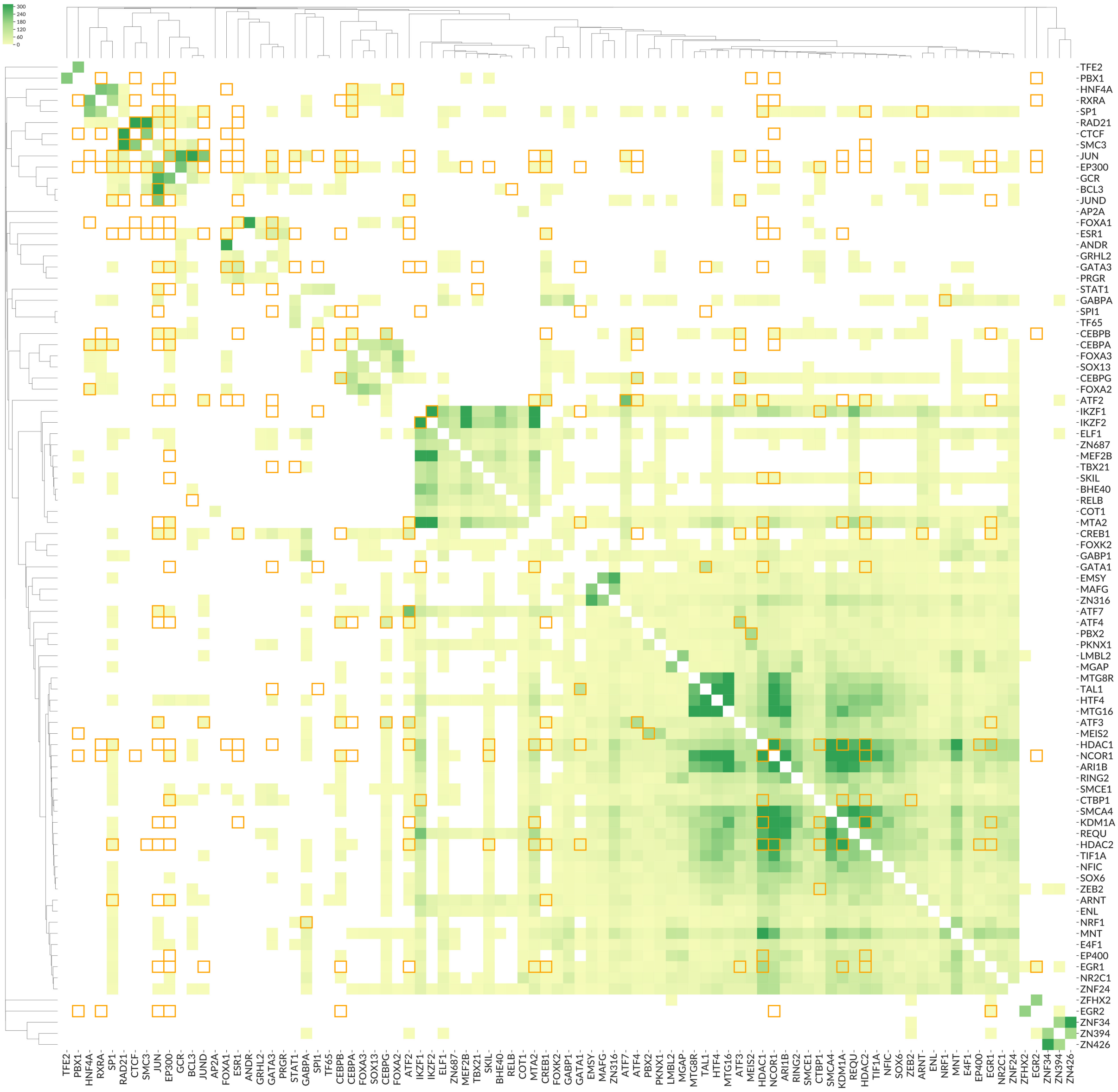

**Supplementary Fig. S7. ASB concordance with binding motifs for different transcription factors.**

Scatterplots (left subpanels) and staveplots (right subpanels) illustrating ASB motif concordance for the top 10 transcription factors with the highest number of ASB-overlapping motif predictions, see **Fig. 4** legend for details. Left column (top to bottom): ANDR, CEBPB, CREB1, CTCF, DUX4. Right column (top to bottom): ESR1, FOXA1, NRF1, SNAI2, SPI1.

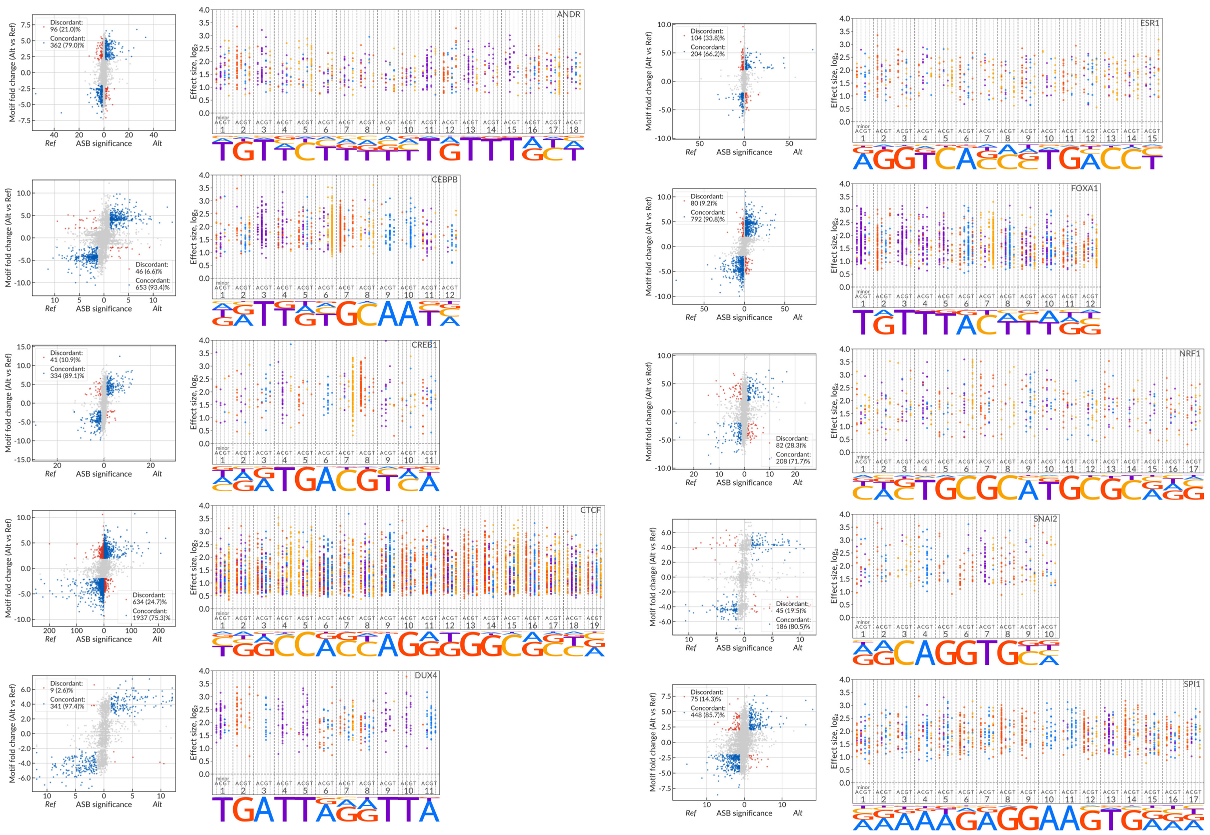

**Supplementary Fig. S8. Performance of the Random Forest classifier for prediction of the TF- and cell type-ASB.**

4 top panels: receiver operating characteristic (ROC) and precision-recall curves for the complete set of ASBs. 12 bottom panels: ROC and PR curves for individual classifiers for 3 transcription factors and cell types with the largest number of ASBs. Area-under-curve values are shown in legends. Transparent curves denote results of individual chromosome hold-outs, bright-colored solid lines show averaged curves.

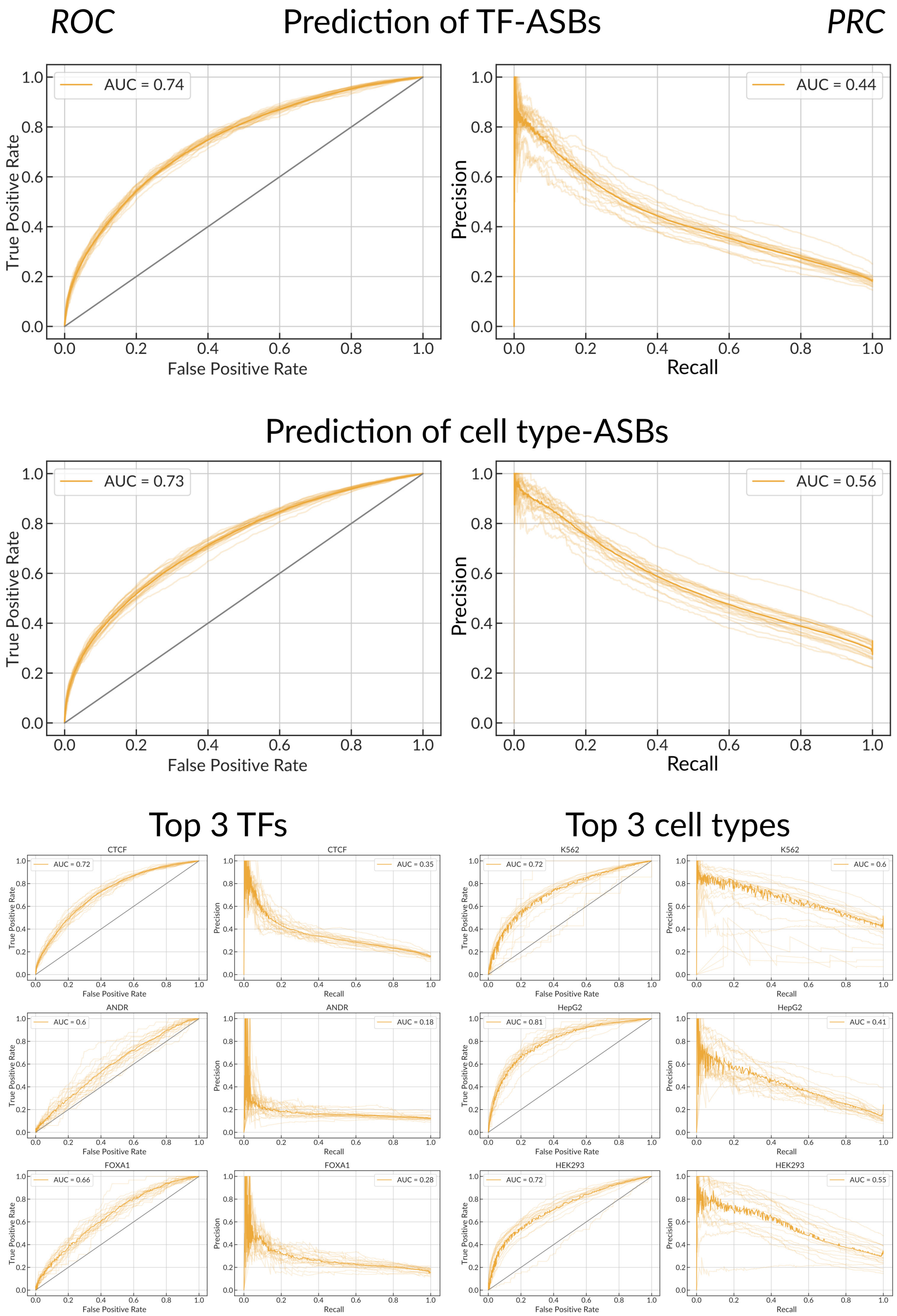

#### Supplementary Fig. S9. ASBs coincide with phenotype-associated SNPs and eQTLs.

**(A-B)** Enrichment of cell type-ASBs among phenotype-associated and eQTL SNVs. Y-axis denotes several mutually exclusive groups of SNPs: CL1 ↑ CL2 ↓ , SNVs carrying both Ref- and Alt-ASBs of different cell types; CL1 ↑ CL2 ↑ , SNVs carrying ASBs for at least two cell types with the same allele preference; single-CL, SNVs with ASB for a single cell type; Low-coverage SNVs that did not pass a total coverage threshold ≥ 20. Non-ASBs are SNVs with the CL-ASB FDR >0.05. X-axis: **(A)** the number of unique (dbSNP ID, trait, database) triples for a given SNV considering four databases of SNP-phenotype associations (EBI, ClinVar, PheWAS, and BROAD autoimmune diseases fine-mapping catalog); **(B)** the number of eQTL target genes according to GTEx eQTL data. The coloring denotes the odds ratios of the one-tailed Fisher's exact test for the enrichment of SNVs with associations for each group of ASBs (against all other SNVs in the table). The gray cells correspond to non-significant enrichments with P > 0.05 after Bonferroni correction for the total number of cells. The values in the cells denote the numbers of SNVs.

**(C-D)** Significant TF-phenotype associations estimated for SNPs with ADASTRA TF-ASBs and EBI-GWAS catalog data. Phenotypes categories: **(C)** cancer, **(D)** immune system disorder. X-axis: TFs, Y-axis: phenotypes. Each bubble represents a TF-phenotype pair with the SNVs found in TF ChIP-Seq data significantly enriched with the phenotype associations (the FDR-corrected P < 0.05 & odds ratio > 2) . The numbers in superscript show the number of TF-ASB sites associated with the phenotype. The area of the circles is proportional to the $\log_2$-number of the phenotype-associated SNVs found in TF ChIP-Seq data. The coloring scheme represents the odds ratios of the enrichment. TF-phenotype combinations w/o ASBs are not shown.

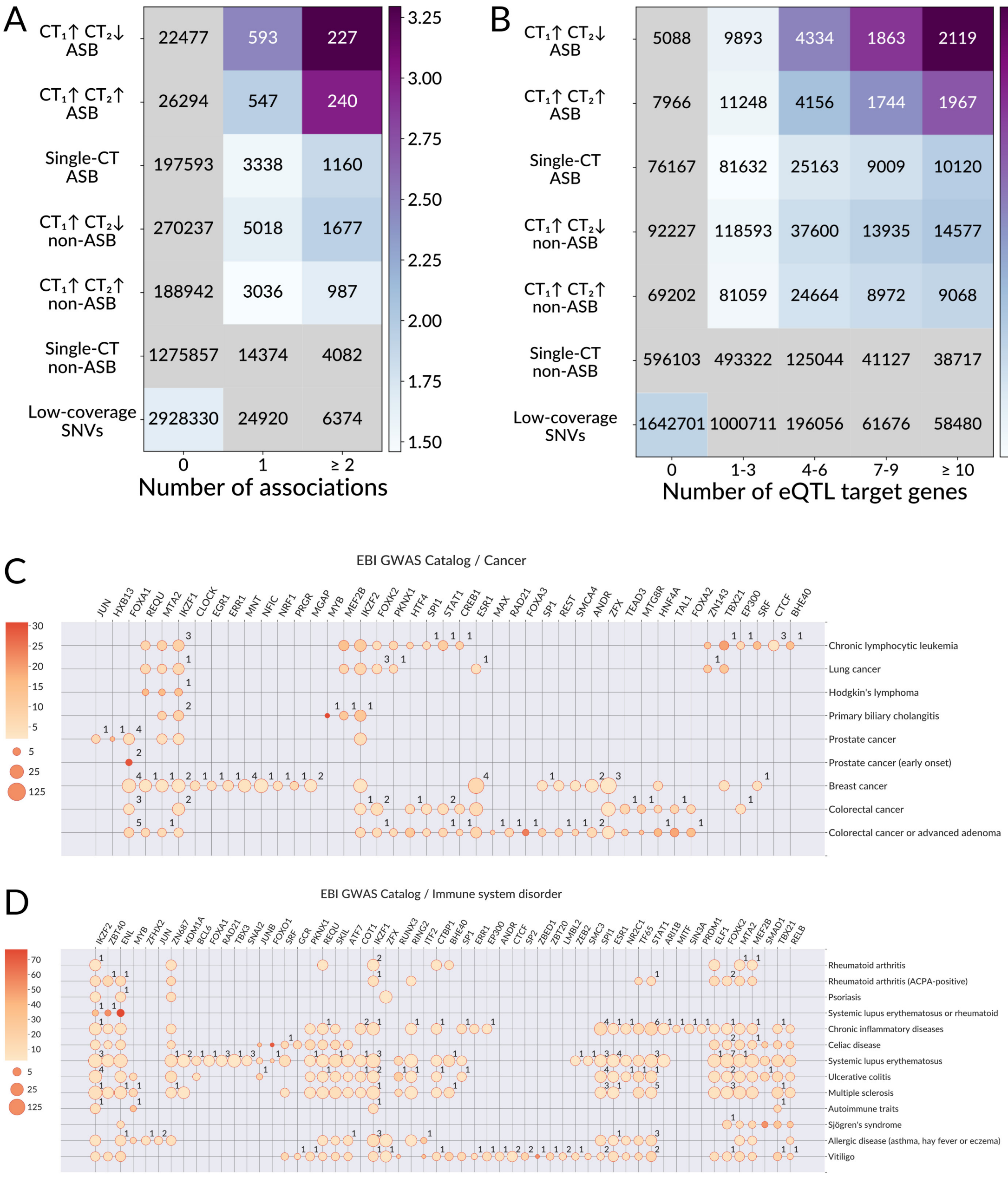

#### Supplementary Fig. S10. Allelic read count distributions at candidate SNVs segregated by BAD.

Rows correspond to different BAD, three panels for each BAD are given (left-to-right):
(left) Heatmap of allelic read count distributions, coloring scheme represents the number of SNVs with the particular read count at the reference allele (X-axis) and the alternative allele (Y-axis);
(middle, right) Distributions of read counts at the reference alleles for SNVs with fixed total coverage denoted by the diagonal slices of the heatmap on the first panels (see the dashed lines in the bottom left corner, the colors correspond to those of the X-axes of the middle and right panels). The purple line represents the non-weighted mixture of Binomial distributions with $p$=1/(BAD+1) and $p$=BAD/(BAD+1), and $n$=total coverage.

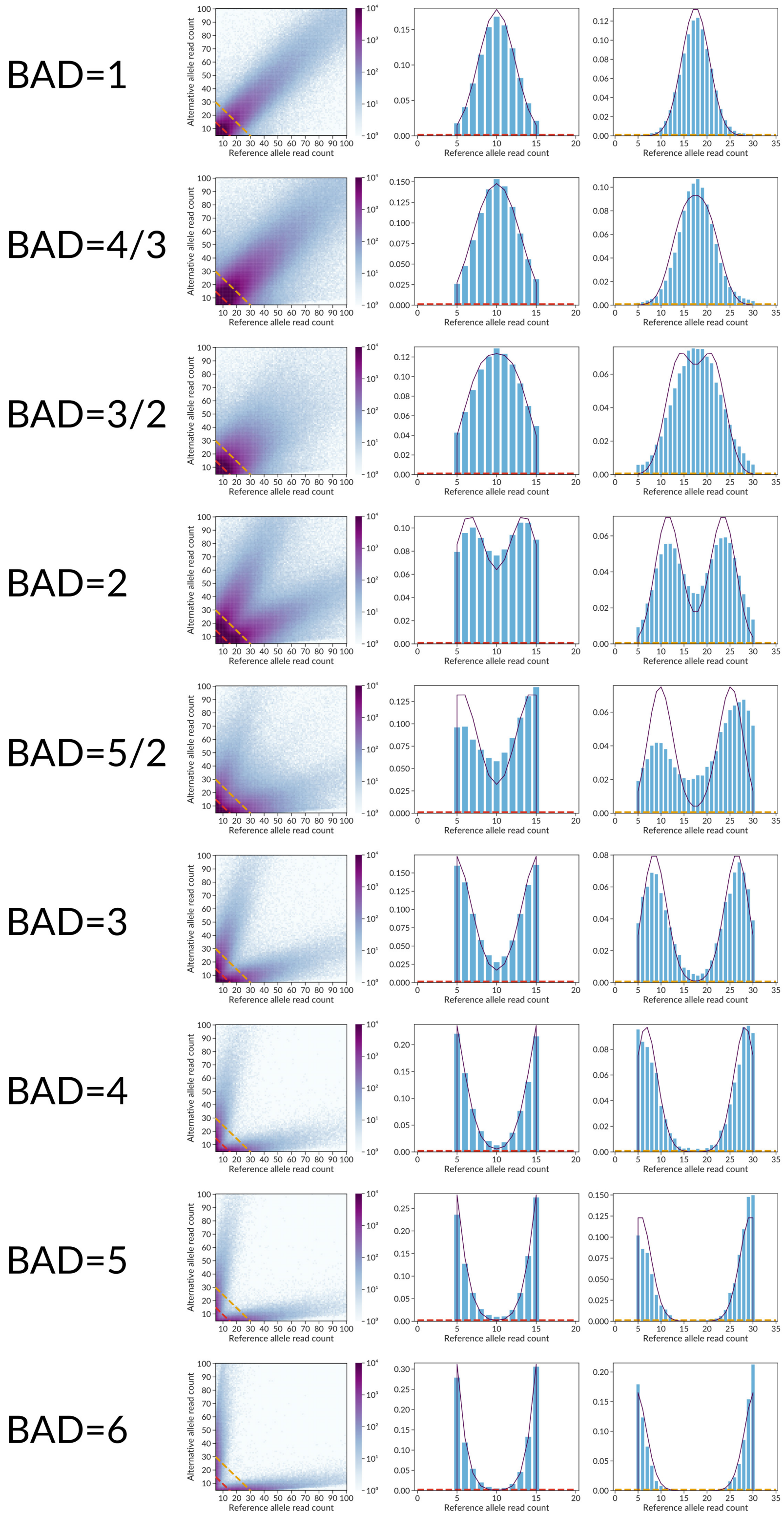

#### Supplementary Fig. S11. The goodness of fit of Negative Binomial mixture models.

The RMSEA (Root Mean Square Error of Approximation) of the fits (Y-axis) plotted against read counts (X-axis) at the reference allele (green) and alternative allele (blue) of the SNVs used for the fits. The fits were performed separately for each BAD. The horizontal dashed line corresponds to RMSEA=0.05 (fits below the threshold were considered non-reliable and the respective SNVs were excluded from ASB calling).

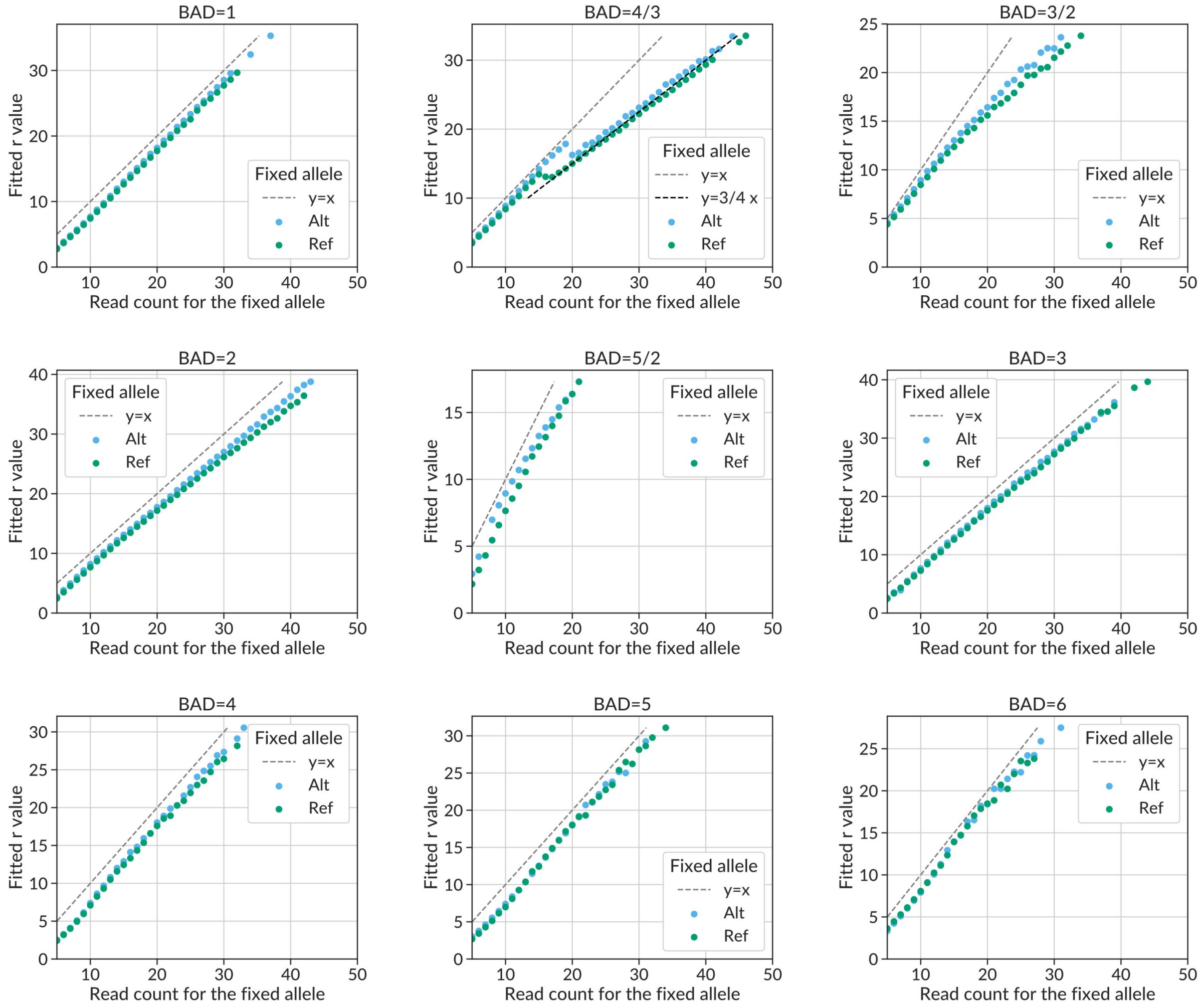

#### Supplementary Fig. S12. The fitted $r$ of the Negative Binomial mixture depending on the allelic read counts.

X-axis: read counts on the reference allele (green) and the alternative allele (blue) of the SNVs that were used for a particular fit. Y-axis: the fitted value of the $r$ parameter. The expected value of $r$ is the number of reads at the fixed allele and is shown as a dashed grey $y=x$ line. For the fixed Alt, distributions of $r$ are systematically skewed to the right due to the reference allele mapping bias; accordingly, the same statistical significance for Ref-ASB is achieved for a higher disbalance value than that for Alt-ASB, which automatically compensates for the mapping bias.

BAD=4/3 has a y=3/4x trend for both fixed reference and fixed alternative read counts, showing that BABACHI provides conservative BAD estimation, with BAD=1 regions being frequently overestimated as BAD=4/3 (see **Supplementary Fig. S3**). The fitted $r$ value for these SNVs, however, compensates for this effect, by fitting the parameters that make the expected value close to that of BAD=1.

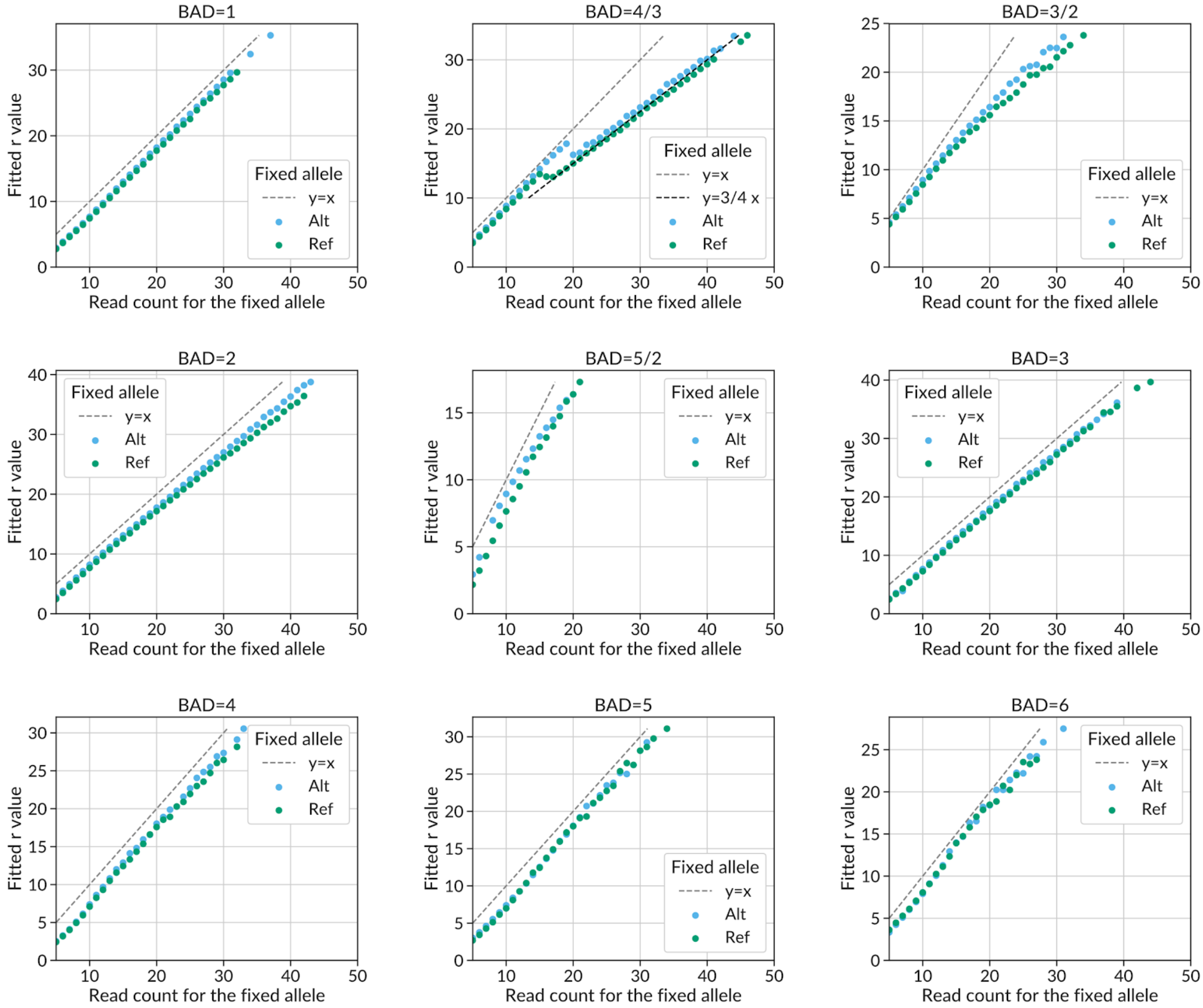

### Supplementary Tables 1-6

#### Table S1. Existing resources providing data on ASBs or chromatin accessibility.

| Type of experiment | Num. of datasets | Num. of SNVs tested for allelic imbalance | Num. of imbalanced SNVs | Num. of cell types | Database name or first author, year | Reference |
|---|---|---|---|---|---|---|
| DNase-Seq | 493 | 362 284 | 64 597 | 121 | Maurano, 2015 | doi:10.1038/ng.3432 |
| TF ChIP-Seq | 177 | 12 490 | 842 | 49 | Maurano, 2015 | doi:10.1038/ng.3432 |
| TF ChIP-Seq | 264 | N/A | 9 962 | 4 | Cavalli, 2016 | doi:10.1007/s00439-016-1654-x |
| ChIP-Seq | N/A | N/A | 17 293 | 7 | Cavalli, 2019 | doi: 10.1038/s41598-019-39633-0 |
| ChIP-Seq | 617 | 298 367 | 14 436 | 3 | Korbolina, 2018 | doi: 10.1002/humu.23425 |
| TF ChIP-Seq | 287 | 135 044 | 7 066 | N/A | AlleleDB, 2016 | doi:10.1038/ncomms11101 |
| TF ChIP-Seq | 45 | 19 106 | 5871 | 8 | Shi, 2016 | doi:10.1093/nar/gkw691 |
| TF ChIP-Seq | >7000 | 2 024 836 | 279 303 | 337 | ADASTRA, 2020 | This study |

#### Table S2. Existing strategies to deal with ploidy, CNVs, and read mapping bias in ASB calling.

| Study | Reference | Cell types | Strategy to deal with the mapping bias | Strategy to deal with CNVs/aneuploidy | Statistical test |
|---|---|---|---|---|---|
| AlleleSeq 2011 | doi:10.1038/msb.2011.54 | GM12878 | Modified binomial test *p* according to mapping probability to each allele (accounts both for reference and inherent bias). | Filtered out CNV regions predicted by read-depth analysis of control gDNA (using CNVnator by Abyzov et. al.). | Binomial test |
| iASeq 2012 | doi:10.1186/1471-2164-13-681 | GM12878 | Reference bias included in the model and estimated using all reads mapped to heterozygous SNPs in each sample. Inherent bias is estimated via simulation. | No adjustment. | Hierarchical Binomial Bayesian model |
| ABC 2015 | doi:10.1093/bioinformatics/btv321 | GM12878 | Reads counts from the control gDNA used to obtain the expected *p* for the binomial test. | Reads counts from the control gDNA used to obtain the expected *p* for the binomial test. | Binomial test |
| Cavalli et. al. 2016 | doi:10.1007/s00439-016-1654-x | GM12878, H1-hESC, K562, SK-N-SH | Reference bias: 2 genomes were reconstructed from haplotypes, alignments to those were obtained separately. Inherent bias: not addressed. | (1) Available phased CNV were used to adjust *p* (only for total copy number less than 3).<br>(2) Filtered out non-diploid regions when only non-phased CNVs were available. | Binomial test |
| AlleleDB 2016 | doi:10.1038/ncomms11101 | 287 ChIP-seq for 14 individuals | Reference bias: reads were aligned to the phased genomes obtained from known phased SNVs with the VCFtodiploid software. Inherent bias: discard reads that had at least one that non-uniquely mapping among all possible mappings. | Filtered out CNV regions, predicted by read-depth analysis of control gDNA (using CNVnator by Abyzov et. al.) | BetaBinomial model |
| Shi et. al. 2016 | doi:10.1093/nar/gkw691 | GM12878 and 6 other GM-, HeLa-S3 | Reference bias: available personalized genomes were used for SNP calling<br>Inherent bias: filtered out sites with notable mapping bias (>0.6) based on the read mapping bias simulation. | Known CNV regions were filtered out. | Binomial test with *p*=0.5 |
| BaalChIP 2017 | doi:10.1186/s13059-017-1165-7 | 14 different cell lines | Reference bias: the average allele ratio in a particular cell type is used as a parameter in a bayesian model.<br>Inherent bias: reads were discarded if mapped incorrectly in the mapping simulation analyzing four possible combinations (ref/alt and +/-). | SNP-wise reference allele frequencies were obtained from external data (B-allele frequencies and phased haplotypes) and then used as parameters in a bayesian model. | BetaBinomial Bayesian model |
| Korbolina et. al. 2018 | doi: 10.1002/humu.23425. | K-562, MCF7, HCT116 and 25 other colon cancer cell lines | Reference bias: alternative genomes were constructed from known SNPs. Inherent bias: not addressed. | No adjustment. | Binomial test with *p*=0.5 |
| Cavalli et. al. 2019 | doi: 10.1038/s41598-019-39633-0 | 7 lGM- cell lines | Identical to Cavalli et. al. 2016 | Known CNV regions from GM12878, centromeric and telomeric regions were filtered out. | Binomial test with *p*=0.5 |

#### Table S3. Quality assessment for BAD predicted from SNP calls for 76 cell types matching between COSMIC and ADASTRA.

| BAD | 2 | 1 | 3 | 3/2 | 4 | 6 | 5/2 | 5 | 4/3 | Other |
|---|---|---|---|---|---|---|---|---|---|---|
| AUROC | 0.849 | 0.835 | 0.866 | 0.871 | 0.492 | 0.997 | 0.393 | 0.462 | 0.635 | - |
| AUPRC | 0.850 | 0.724 | 0.665 | 0.340 | 0.016 | 0.787 | 0.002 | 0.003 | 0.000 | - |
| Precision* | 0.886 | 0.866 | 0.856 | 0.240 | 0.067 | 0.660 | 0.001 | 0.008 | 0.240 | - |
| Recall* | 0.698 | 0.166 | 0.505 | 0.784 | 0.031 | 0.882 | 0.011 | 0.007 | 0.784 | - |
| *% of total SNPs* | *45.89* | *36.99* | *8.94* | *5.99* | *0.76* | *0.22* | *0.21* | *0.170* | *0.04* | *0.78* |
| *% of total, cumulative* | *45.89* | *82.88* | *91.82* | *97.81* | *98.57* | *98.79* | *99.00* | *99.17* | *99.21* | *100.00* |

*Precision and Recall values are computed by using the BAD maps as the multiclass classifiers.

#### Table S4. Features used for applying machine learning to predict ASBs.

| Feature type | Feature ID | Description |
|---|---|---|
| ADASTRA SNV annotation Motif-based | motif_log_pref, motif_log_palt | $-\log_{10}$(P-value) of the motif hit for the Ref and Alt alleles |
| | motif_fc | Log-ratio of P-values for motif hits at the Ref and Alt alleles |
| | motif_pos | Relative position of the SNV within the motif occurrence |
| | is_repeat | Whether or not an SNV is located in a DNA repeat region |
| Allele-specific chromatin DNase accessibility based on Maurano et al. data | numhets_dnase | The number of heterozygous samples for a given allele |
| | reads1_dnase, reads2_dnase | Numbers of DNase-seq reads supporting Ref and Alt allelic variants |
| | totalReads_dnase pctRef_dnase | The total number of DNase-seq reads covering an SNP and the fraction of reads supporting Ref allele |
| | qvalue_dnase | Q-value of the allele-specific chromatin accessibility |
| | is_imbalanced_dnase_bool | Whether an SNP passes the FDR 5% to be considered imbalanced |
| Features produced by sequence analysis with DeepSEA artificial neural network | 919 features for Ref allele | 919 neuron outputs of the last DeepSEA layer for the reference allele |
| | 919 features for Alt allele | 919 outputs for the alternative allele |
| | 919 differential features | 919 values of the differences between neuron outputs for the reference and alternative alleles |
| | 919 E-values | E-values for the observed differences |
| | 919 Odds-ratios | Odd-ratios for the observed differences |

**Table S5. Overview of the data used for machine learning of ASBs and results of the model validation.**

| **Top 10 TFs factors used for ASB analysis with machine learning** | | | | |
|---|---|---|---|---|
| **TF** | **TF-ASB** | **Total num. of SNVs** | **AUROC, ind. model** | **AUPRC, ind. model** |
| ANDR | 2016 | 16212 | 0,60 | 0,18 |
| CEBPB | 960 | 12220 | 0,68 | 0,25 |
| CTCF | 8549 | 53950 | 0,72 | 0,35 |
| ESR1 | 3091 | 32668 | 0,66 | 0,17 |
| FOXA1 | 2366 | 14759 | 0,66 | 0,28 |
| FOXK2 | 2840 | 20865 | 0,60 | 0,18 |
| IKZF1 | 1716 | 26616 | 0,64 | 0,13 |
| RAD21 | 1956 | 21818 | 0,72 | 0,21 |
| SPI1 | 2324 | 16731 | 0,62 | 0,21 |
| STAT1 | 2670 | 20298 | 0,68 | 0,24 |
| ***Complete data (All TFs)*** | 42224 | 231355 | | |
| **Top 10 cell types used for ASB classification task** | | | | |
| **Cell type** | **CL-ASB** | **Total num. of SNV.** | **AUROC, ind. model** | **AUPRC, ind. model** |
| A549 lung carcinoma | 4142 | 11619 | 0,76 | 0,64 |
| CD14+ monocytes | 2739 | 20070 | 0,68 | 0,26 |
| GM12878 female B-cells | 11749 | 38067 | 0,68 | 0,47 |
| HEK293 embryonic kidney | 11356 | 37863 | 0,72 | 0,55 |
| HepG2 hepatoblastoma | 4442 | 34575 | 0,81 | 0,41 |
| K562 myelogenous leukemia | 17105 | 40956 | 0,72 | 0,60 |
| LNCaP prostate carcinoma | 1522 | 12967 | 0,67 | 0,27 |
| MCF7 Invasive ductal breast carcinoma | 6241 | 81099 | 0,74 | 0,22 |
| VCaP prostate carcinoma | 1849 | 10605 | 0,69 | 0,33 |
| foreskin keratinocyte | 5062 | 19387 | 0,67 | 0,39 |
| ***Complete data (All cell types)*** | 67824 | 231355 | | |

### Supplementary Methods

Chromosome segmental duplications and aneuploidy results in varying doses of specific allele in different cell types or individuals. This can bias estimation of allelic frequencies in various genetics problems. We introduce the Background Allelic Dosage (BAD) as the ratio of the major to minor allele dosage in the particular genomic segment, which depends on chromosome structural variants and aneuploidy. We have developed a Bayesian Changepoint Identification algorithm inspired by V. E. Ramensky et al.[doi: 10.1089/10665270050081487] and Pasio to construct full-genome BAD maps. It consists of two stages: first, we find long deletions and centromeric regions by comparing distances between neighbouring SNVs; then we use the SNV coverage to predict BAD.

#### 1. Stage 1 of the segmentation

The first stage of the procedure is segmentation by genome positions of SNVs, without taking into account the ratio of read at alternative alleles. The goal is to find centromeric regions, large deletions and other regions depleted of SNVs.

The essence of the Stage 1 algorithm is an iterative recalculation of the effective chromosome length, at each step finding long regions with no SNVs.

Suppose the genome positions of the called SNVs are $P = [s_1, s_2, \ldots, s_N]$. Then the output of the stage 1 algorithm is the array of sub-chromosome boundaries $B = [n_1, n_2, \ldots, n_m], 1 < n_i < N$ - ordinal numbers of SNVs in the original chromosome - such that the spaces between each $s_{n_i}$ and $s_{n_i+1}$ are marked as deletions. These boundaries divide the original chromosome into $m + 1$ sub-chromosomes: $\{[s_1, \ldots, s_{n_1}], [s_{n_1+1}, \ldots, s_{n_2}], \ldots, [s_{n_m+1}, \ldots, s_N]\}$.

At each iteration step the distance between neighbouring SNVs is considered to be a deletion if it exceeds the "critical gap" of $L_{\text{eff}} \cdot (1 - 10^{-\frac{1}{\sqrt{N}}})$, where $L_{\text{eff}}$ is the length of the chromosome in bp excluding all currently found deletions. The intuition behind this formula is as follows: if the SNV positions had uniform distribution along the chromosome, then the expected number of SNVs on any region with the length less than $L_{\text{eff}} \cdot (1 - N^{-\frac{1}{N}})$ would be less than 1. As far as the actual distribution of SNV positions is significantly non-uniform, we use the heuristic factor of $1 - 10^{-\frac{1}{\sqrt{N}}}$, which decreases with N at a slower rate than $1 - N^{-\frac{1}{N}}$ does.

**Algorithm 1** Segmentation by SNVs' positions (Stage 1)

$P \leftarrow [s_1, s_2, \ldots, s_N]$ ▷ The array of SNVs' genome positions

$Eff\_len \leftarrow s_N - s_1$ ▷ Effective length = difference between first and last SNV positions

$CGF \leftarrow 1 - 10^{-\frac{1}{\sqrt{N}}}$ ▷ The critical gap factor

$B \leftarrow \{\}$ ▷ The set of subchromosome borders

**do**

  $Eff\_len \leftarrow Eff\_len - \Delta len$ ▷ Reduce the effective length by the sum of deletions

  $\Delta len = 0$ ▷ Set the sum of all deletions to 0

  **for** $i \leftarrow 1$ **to** $N - 1$ **do**

    **if** $i$ **not in** $B$ **then** ▷ If current SNV is not already in the set of boundaries

      $Dif \leftarrow P[i+1] - P[i]$ ▷ The distance between $i$-th and $i+1$-st SNV

      **if** $Dif > (Eff\_len - Dif) \cdot CGF$ **then** ▷ If the distance exceeds the critical gap

        $\Delta len \leftarrow \Delta len + Dif$ ▷ Add the distance to the sum of deletions

        **add** $i$ **to** $B$ ▷ Add current SNV to the set of boundaries

      **end if**

    **end if**

  **end for**

**while** $\Delta len > 0$ ▷ Repeat while any boundaries are found

**return** $B$

#### 2. Stage 2 of the segmentation

The objective of the second stage of the procedure is to find the regions of approximately constant background allelic dosage (BAD). The value of BAD does not depend on phasing of particular SNVs and thus can be estimated even when haplotype phasing data is not available.

In this stage we first call changepoints (or boundaries) with the marginal likelihood segmentation (inspired by pasio) and then use maximum posterior estimation to assign BAD to every called region.

Below we explain the BAD estimation (Stage 2b) first and the changepoint calling (Stage 2a) last, as far as the same likelihood expression is involved.

##### 2.1 *Stage 2b: BAD estimation*

We assume that allele read counts have binomial distribution, if there is no SNV-specific bias, such as ASB or mapping bias (null hypothesis). With that, the probability to observe $k$ reference read counts ($\mathrm{Ref_c}$) given the total of $n$ read counts on both allele is given by:

$$p(\mathrm{Ref_c} = k) = \binom{n}{k} p^k (1-p)^{n-k} \,. \tag{2.1}$$

The parameter $p$ has the meaning of background allele ratio and is to be estimated. However, to allow for missing haplotype phasing we introduce the following statistics:

$$X = \min(\mathrm{Ref_c}, \mathrm{Alt_c}) \,, \tag{2.2}$$

which, under the binomial distribution assumption, has the following probability distribution:

$$p(X = k) = \begin{cases} \binom{n}{k} p^k (1-p)^{n-k} + \binom{n}{n-k} p^{n-k} (1-p)^k, & \text{if } k < \frac{n}{2} \,, \\ \binom{n}{k} p^k (1-p)^{n-k}, & \text{otherwise } (k = \frac{n}{2}\ ) \,, \end{cases} \tag{2.3}$$

that can be rewritten in a more compact form:

$$p(X=k) = \binom{n}{k} p^k (1-p)^{n-k} \left(1 + \left(\frac{1-p}{p}\right)^{-(n-2k)}\right) \cdot 2^{(-\delta_{n,2k})} \,. \tag{2.4}$$

Let $x_i$ denote the value of the statistics $X$ on the $i$th SNV. With that, the log likelihood to observe $x_i$ is given by $\mathcal{L}_i$ :

$$\mathcal{L}_i(p|x_i,n_i) = \tilde{\mathcal{L}}_i(p|x_i,n_i) + g(x_i,n_i) \,, \tag{2.5}$$

where

$$\tilde{\mathcal{L}}_i(p|x_i,n_i) = x_i \ln p + (n_i - x_i) \ln(1-p) + \ln\left(1 + \left(\frac{1}{p} - 1\right)^{-(n_i - 2x_i)}\right) - \ln(1 - \sigma_i(p_i,n_i)) \,, \tag{2.6}$$

$$g(x_i,n_i) = \ln \binom{n_i}{x_i} - \delta_{n_i,2x_i} \cdot \ln 2 \,, \tag{2.7}$$

and

$$\sigma_i(p_i,n_i) = \sum_{k=0}^{4} p(\xi = k) \,. \tag{2.8}$$

As we filter out the low coverage SNVs, using the filter $Ref_c + Alt_c \geqslant 5$, the likelihood must be corrected for the truncated values of X. This is made by introducing the last term in 2.5 (containing $\sigma_i$ as in 2.7).

The log likelihood of a region with $N$ SNVs is the sum of SNV-wise likelihoods (assuming the values of the statistics $x$ on all SNVs to be independent):

$$\mathcal{L}(p) = \sum_{i=1}^{N} \mathcal{L}_i(p) \,. \tag{2.9}$$

As far as the above likelihood is symmetrical with respect to the transformation $p \to (1-p)$ (interchange of the alleles), we limit our estimation to $p \in [0; 0.5]$ and define BAD on the segment to be $\frac{1}{1+\hat{p}}$. Moreover, as far as cells rarely have copy numbers higher than 7 we further limit the estimation to discrete set of allele ratios $\Omega = \{\frac{1}{2}, \frac{3}{7}, \frac{2}{5}, \frac{1}{3}, \frac{2}{7}, \frac{1}{4}, \frac{1}{5}, \frac{1}{6}, \frac{1}{7}\}$, corresponding to $TotalCN \leqslant 7$. This allows one to define the estimate of the parameter $\hat{p}$ on a region with SNVs to be the maximum a posteriori estimation:

$$\hat{p} = \arg\max_{p \in \Omega} \left(\mathcal{L}(p) + \log \pi(p)\right) , \tag{2.10}$$

where we consider

$$\pi(p) = \frac{1}{|\Omega|} , \tag{2.11}$$

- to be an uninformative prior.

#### 2.2 *Stage 2a: Marginal Likelihood Segmentation*

To find the changepoints (and therefore the regions of consistent BAD) we use the principal of extreme marginal likelihood. The marginal likelihood of a region with no changepoints is the probability to observe the series of $x_i$ on the SNVs of the region assuming independent binomial trials with some common $p \in \Omega$. The logarithmic likelihood is given by:

$$L = \log\left[\sum_{p \in \Omega} \pi(p) \exp\left(\mathcal{L}(p)\right)\right] = \log\left[\sum_{p \in \Omega} \pi(p) \exp\left(\sum_{i=0}^{N} \mathcal{L}_i(p)\right)\right] . \tag{2.12}$$

For a region with $M-1$ changepoints (that is $M$ segments), its log marginal likelihood is the sum of log marginal likelihoods of its segments.

$$\mathscr{L} = \sum_{k=1}^{M} L^k , \tag{2.13}$$

where

$$L^k = \log\left[\sum_{p \in \Omega} \exp\left(\sum_{i=1}^{N_k} \mathcal{L}_i^k(p)\right)\right] , \tag{2.14}$$

is the log marginal likelihood of the $k$-th segment and $N_k$ is the number of SNVs it contains.

Finally, we use the CAIC (Consistent Akaike Information Criterion) to compare likelihoods of segmentations with different number of changepoints. This is equivalent to adding $\Delta_{boundaries}$ to the log marginal likelihood

$$\Delta_{\text{boundaries}}(b) = -\frac{|\Omega| \cdot b}{2}(1 + \log N)\,, \tag{2.15}$$

where $b = M - 1$ is the number of boundaries of a particular segmentation and $|\Omega| = 9$ (the number of allowed states). Finally, the expression of log marginal likelihood of the segmentation is as follows:

$$\mathscr{L} = \sum_{k=1}^{M} \log\left[\sum_{p\in\Omega} \exp\left(\sum_{i=1}^{N_k} \tilde{\mathcal{L}}_i^k(p)\right)\right] + \Delta_{\text{boundaries}}(M-1) - \log|\Omega| + \sum_{j=1}^{N} g(x_i, n_i)\,. \tag{2.16}$$

We find the changepoints on the subchromosome that maximize $\mathscr{L}$ with the help of dynamic programming algorithm.

#### 2.3 *Dynamic programming algorithm for Stage 2a*

For a subchromosome with $N$ SNVs we define a segmentation to be a set of boundaries $\{S = k_i\}$ where $k_i$ is the ordinal number of the SNV previous to $i$-th boundary,
$0 = k_0 \leqslant k_1 \leqslant \ldots \leqslant k_b \leqslant k_{b+1} = N$ ($b$ is the number of added boundaries). Let $S_m$ be the optimal segmentation of the first $m$ SNVs of the subchromosome (as if there are no other SNVs), $b_m$ be the number of added boundaries of that segmentation and let $W[k,l]$ be the marginal logarithmic likelihood of a segment of the subchromosome from $k$-th to $l$-th SNV

$$W[k,l] = \log\left[\sum_{p\in\Omega} \exp\left(\sum_{i=k}^{l} \tilde{\mathcal{L}}_i(p)\right)\right]. \tag{2.17}$$

(here we omitted the boundaries term and the terms that do not depend on S)

Then we define the score of a segmentation to be:

$$\text{score}(S) = \sum_{i=0}^{M-1} W[k_i + 1, k_{i+1}] + \Delta_{\text{boundaries}}(M-1)\,, \tag{2.18}$$

The segmentation with extreme score also maximizes $\mathscr{L}$ and therefore is the optimal segmentation.

With that score($S_{m+1}$) can be found from score($S_m$) by the following recurrent expression:

$$\begin{cases} \text{score}(S_{j+1}) = \max\limits_{k\in[0;j]} (\text{score}(S_k) + \Delta_{\text{boundaries}}(b_k+1) - \Delta_{\text{boundaries}}(b_k) + W[k+1;j+1]) \\ k_{max} = \arg\max\limits_{k\in[0;j]} (\text{score}(S_k) + \Delta_{\text{boundaries}}(b_k+1) - \Delta_{\text{boundaries}}(b_k) + W[k+1;j+1]) \\ b_{j+1} = b_{k_{max}} + 1 \\ b_0 = \text{score}(S_0) = 0 \end{cases} \tag{2.19}$$

The algorithm has O($|\Omega|N^2$) time complexity.

#### 2.4 *Dynamic programming optimization*

The above procedure was modified to achieve O($|\Omega|NlogN$) time complexity.

The array of SNVs of the subchromosome is divided into pieces with approximately 600 SNVs in each with overlap of 300 SNVs. On each piece the original segmentation stage 2 algorithm was performed to get the set of candidate boundaries of the piece. Then the candidate boundaries sets from all pieces were united to get the candidate boundaries set for the subchromosome. Finally, the stage 2 algorithm was applied to the whole subchromosome, only allowing boundaries placement among the candidate boundaries. This was made by summing up likelihoods of SNVs inside regions between every consequent candidate boundaries pair and treating these SNVs as a single object (SNV).

SNVs on the same position (that came from aggregation of multiple datasets) were also treated as one SNV with the likelihood of sum of such SNVs.
